# Changes in wheat rhizosphere microbiota in response to chemical inputs, plant genotype and phenotypic plasticity

**DOI:** 10.1101/2021.05.07.441152

**Authors:** Samuel Jacquiod, Tiffany Raynaud, Eric Pimet, Chantal Ducourtieux, Leonardo Casieri, Daniel Wipf, Manuel Blouin

**Affiliations:** Agroécologie, AgroSup Dijon, INRAE, Univ. Bourgogne, Univ. Bourgogne Franche-Comté, F-21000 Dijon, France; Agroécologie, AgroSup Dijon, CNRS, INRAE, Univ. Bourgogne, Univ. Bourgogne Franche-Comté, F-21000 Dijon, France; Mycorrhizal Applications LLC, c/o BRDG Park at the Danforth Center. St. Louis, Missouri (USA)

**Author notes:** Corresponding author: Manuel Blouin.

**Keywords:** ancient varieties, bacteria, fungi, genotype, inputs, landraces, microbial community, modern varieties, mycorrhiza, phenotypic plasticity, rhizosphere, wheat

## Abstract

Since modern wheat varieties are grown with chemical inputs, we ignore if changes observed in rhizosphere microorganisms between ancient and modern varieties are due to i) breeding-induced changes in plant genotype, ii) modifications of the environment via synthetic chemical inputs, or (iii) phenotypic plasticity, defined as the interaction between the genotype and the environment. In the field, we evaluated the effects of various wheat varieties (modern and ancient) grown with or without chemical inputs (N-fertilizer, fungicide and herbicide together) in a crossed factorial design. We analysed rhizosphere bacteria and fungi by amplicons sequencing and mycorrhizal association by microscopic observations. When considered independently of plant genotype, chemical inputs were responsible for an increase in dominance for bacteria and decrease in evenness for bacteria and fungi. Independently of inputs, modern varieties had richer and more even bacterial communities compared to ancient varieties. Phenotypic plasticity had a significant effect: bacterial and fungal diversity decreased when inputs were applied in ancient varieties but not in modern ones. Mycorrhiza were more abundant in modern than ancient varieties, and less abundant when using chemical inputs. Although neglected, phenotypic plasticity is important to understand the evolution of plant-microbiota associations and a relevant target in breeding programs.

## INTRODUCITON

Rhizosphere microbiota is involved in several key functions for the plant, such as the regulation of nutrient access, environmental disturbance tolerance and disease resistance (Rodriguez et al., 2008; Mei & Flinn, 2010; Farrar et al., 2014). There is abundant evidence that host genotypes have a significant influence on rhizosphere microbial community composition (Lundberg et al., 2012; Peiffer et al., 2013) and its function, with consequences on plant growth, development and immunity (Lemanceau et al., 2017). In agriculture, understanding factors that favour crop’s associations with beneficial bacteria and fungi could help in maintaining high crop yields without using synthetic chemical inputs. Breeding and application of synthetic chemical inputs are two interconnected agricultural practices, which can modify plant-microbiota interactions. Therefore, studying the effect of domestication on the rhizosphere microbiota without considering chemical inputs does not allow an accurate understanding of the evolution of plant-microbiota interactions. The effects of breeding and inputs can be described by adopting the formalism of quantitative genetics P = G + E + G×E (Falconer, 1989). In our case, the plant-microbiota interactions can be considered as a phenotypical trait “P”, which can be determined by: (i) the plant genotype “G” (either modern or ancient varieties; microorganisms genotypes will not be considered explicitly here), (ii) the environment “E” (modified by agricultural practices such as the application of inputs) and (iii) the interaction between crop genotype and inputs “GxE” (defined as plant phenotypic plasticity). Indeed, previously reported observations call for considering these three important drivers of plant-microbiota interactions, which we will briefly present in the following order: E, G and G×E.

First, synthetic chemical inputs applied in the fields since the Green Revolution can be considered as a modification of the environment (E), with direct modifications of the soil and rhizosphere microbial communities. In a meta-analysis, long term mineral fertilizer application has been shown to increase microbial biomass by 15.1% (Geisseler and Scow, 2014). Fertilization can change microbial community composition, by promoting copiotroph organisms such as specific members from Actinobacteria and Firmicutes, whereas decreasing the oligotroph organisms such as specific members from Acidobacteria and Verrucomicrobia (Ramirez et al. 2012). These modifications of the soil microbial communities can reverberate on the rhizosphere community. For example, nitrogen application selects less mutualistic rhizobia, with less benefit to the host (Weese et al. 2015). In a long-term experiment, Ai et al. (2015) demonstrated that inorganic fertilization decreased the rhizospheric dependence on root derived carbon for Actinobacteria members. Similarly, plants-AMF (arbuscular mycorrhizal fungi) symbiotic association can also be altered by inorganic fertilization (Lamber et al., 2009), with changes in the diversity of AMF (EgertonLWarburton et al., 2007).

Second, some changes in plant-microbiota interactions can arise from modifications in the plant genome (G), through mutation, hybridization or allele fixation. Domesticated plants changed their interactions with soil microorganisms compared to their wild relatives (Garcia-Palacios et al., 2013; Milla et al., 2015; Turcotte et al., 2015). Artificial selection for improved yield in high-input agriculture could unintentionally lead to a reduction of root microbiota members involved in nutrient acquisition or plant immunity under low input conditions (Perez-Jaramillo et al., 2016). The effect of domestication (from wild relatives to cultivated crops) on rhizosphere microbiota has been observed in several crops such as barley (Bulgarelli et al., 2015), maize (Szoboszlay et al. 2015), foxtail millet (Chaluvadi and Bennetzen, 2018) and common bean (Perez-Jaramillo et al., 2017). In wheat, the rhizosphere bacterial communities of ancient varieties was more diverse than modern varieties (Germida and Siciliano, 2001). The general pattern is a rhizospheric enrichment of Actinobacteria and Proteobacteria members and a decrease in Bacteroidetes members in modern varieties compared to wild relatives (Perez-Jaramillo et al., 2018). Changes in plant-microbes interactions between wild relatives, ancient varieties and modern varieties were also observed for mycorrhizal associations. Many studies showed that the mycorrhizal association and responsiveness of modern wheat varieties in terms of yield gain was lower than that of the ancient varieties or wild relatives (Kapulnik and Kushnir 1991; Hetrick et al., 1992; Zhu et al., 2001; Leiser et al., 2016). However, a meta-analysis showed that modern varieties were less intensely colonized but were more mycorrhiza-responsive compared to ancestral genotypes, concluding on the absence of evidence that agricultural and breeding practices are responsible for a lack of response to mycorrhiza in new crop genotypes (Lehmann et al., 2012).

Third, artificial selection may have influenced the way plant genotypes respond to inputs in terms of plant-microbiota interaction (G×E), i.e. the phenotypic plasticity of plant-microbiota interaction. Phenotypic plasticity denotes the ability of a genotype to exhibit changes in a specific trait across different environments (Laitinen et al., 2019). Artificial selection of modern varieties has been very efficient in providing agriculture with productive cultivars displaying stable performances across diverse environmental conditions, but it is not clear if these stable yields are linked to the phenotypic plasticity of other traits (Gage et al., 2017), such as those involved in plant-microbiota interaction. Several authors thus suggest that understanding G×E interactions could be a novel breeding strategy, coping better with changing environments while securing stable yields. Since genes responsible for yield (i.e. mean trait value) and phenotypic plasticity (i.e. variance) are distinct, breeders should theoretically be able to select both at the same time to generate plastic varieties that adapt better to a changing environment, while maintaining a decent yield (Kusmec et al., 2017).

Therefore, it is necessary to evaluate the relative importance of E, G and the G×E interaction for a more accurate understanding of the plant-microbiota interactions in the rhizosphere of selected crops. Specifically, evaluating the contribution of these parameters can help to determine if artificial selection has played an important role in the evolution of plant-microbiota interactions through genetic effects, being either independent (G) or dependent (G×E) of the environment. In this aim, we studied the structure of rhizosphere microbial community of ancient and modern varieties of wheat, in the presence or absence of inputs (N-fertilizer, fungicide and herbicide). We hypothesized that i) the presence of inputs decreases the microbial diversity in the rhizosphere, ii) modern genotypes have lost their ability to establish interactions with some microbial species, iii) the lower plant-microbiota association in modern genotypes is amplified in the presence of inputs. We used an integrated approach coupling the analysis of microscopic observations of mycorrhiza, and amplicon sequencing of the bacterial (16S rRNA gene) and fungal (ITS2) communities.

## MATERIALS AND METHODS

### Field site description

The field experiment was conducted on dedicated plots at AgroSup Dijon, the Institut National Supérieur des Sciences Agronomiques de l’Aliment et de l’Environnement (47 ° 18’32 “N 5 ° 04’02” E, Dijon, France). The climate of the experimental area is semi-continental, with an average annual temperature of 11 ^°^C (±4.5^°^C). Average precipitation per year is 760.5 mm, monthly sunshine hours are 1848.8 h. The clay loam soil characteristics in the 0-22 cm horizon were as follow: pH_H2O_ = 8.0; 34.6% clay, 36.2% loam, 29.2% sand; 26.7 g.kg^-1^ organic carbon (46.2 g.kg^-1^ organic matter), 2.11 g.kg^-1^ total nitrogen; 294.0 g.kg^-1^ total Ca; 0.020 g.kg^-1^ P_2_O_5_; 24.30 cmol^+^.kg^-1^. The preceding culture was a field bean (*Vicia faba*) for all the experimental plots.

### Experimental design and sampling

Two kinds of genotypes, hereafter called “breeding types” consisting of five modern and five ancient wheat varieties were sown in the 3^th^ and 4^th^ of November 2016. Modern varieties were selected after the 60’s in agrosystems concomitantly receiving high levels of inputs: Rubisko (R, 2012) provided by RAGT Semences, Descartes (D, 2014) and Sherlock (S, 2015) provided by Secobra, Alixan (A, 2005) provided by Limagrain and Tulip (T, 2011) provided by Saaten Union (http://www.fiches.arvalis-infos.fr/). Ancient varieties were provided by the non-governmental organization “Graines de Noé” (http://www.graines-de-noe.org/), which promotes the conservation of wheat landraces. Among their 200 varieties, all grown without inputs, we selected some with a local origin: Barbu du Mâconnais (BM, XIXth–beginning XXth century), Blé de la Saône (BS, before 1960), Automne Rouge (AR, XIXth century). To diversify the panel of wheat grown without synthetic inputs, we also selected Alauda (AL), a variety dedicated for organic agriculture or biodynamic, obtained in 2013 by crossing the varieties Probus (1948) and Inntaler (before 1960) and einkorn wheat (*Triticum monococcum*, EW) which is increasingly grown by farmers interested in ancient varieties (https://urgi.versailles.inra.fr/Projects/Achieved-projects/Siregal). Two growing conditions were tested for each variety: i) with inputs (w) and ii) without inputs (w/o), with three replicates of each condition distributed in three blocks. Inputs included fertilizer (CAN 27% Granulé, Dijon Céréales, France) for a total of 150 kg N.ha^-1^, applied as 50 kg N.ha^-1^ in three times, on week 18 after sowing (09/03/2017), week 25 (24/04/2017) and week 30 (30/05/2017)); herbicide (Bofix™, Dow Agro Science, supplied once at 0.3 l.ha^-1^ on week 23, the 10/04/2017); and fungicide (Bell Star™, Dow Agro Science), applied once at 2.5 kg.ha^-1^ on week 26, the 05/05/2017. The experiment thus consisted in 60 plots (five modern and five ancient varieties, with or without inputs, replicated three times) of 1m² each, separated from the other by 0.8m. In each plot, 300 seeds of each variety were sown manually in seven rows.

### Sampling of rhizosphere microbial communities

In each plot, we sampled randomly two wheat rhizospheres. After loosening the soil with a fork, the root system of the plant with its root-adhering soil was extracted from the bulk soil. In the laboratory, rhizosphere soil was gently removed from the roots by brushing. The two samples from the same plot were pooled together to obtain a representative sample. All samples were frozen at −20°C until further processing for DNA extraction.

### Microbial community analysis

Total bacterial and fungal diversity and composition from rhizosphere soil samples were respectively analyzed by sequencing the 16S rRNA gene V3-V4 region, and the ITS2 region via Illumina Miseq 2x 250 bp paired-end analysis. Total DNA was extracted from 250 mg of rhizospheric soil using the DNeasy PowerSoil-htp 96 well DNA isolation kit (Qiagen, France). In two steps, 16S rRNA gene and ITS2 amplicons were generated for all extracts. In the first step, the V3-V4 hypervariable region of the bacterial 16S rRNA gene was amplified by polymerase chain reaction (PCR) using the fusion primers U341F (5’-CCTACGGGRSGCAGCAG-3’) and 805R (5’-GACTACCAGGGTATCTAAT-3’), with overhang adapters (forward: TCGTCGGCAGCGTCAGATGTGTATAAGAGACAG, reverse: GTCTCGTGGGCTCGGAGATGTGTATAAGAGACAG) to allow the successive addition of multiplexing index-sequences. Fungal ITS2 was amplified using the primers ITS3F (5’-GCATCGATGAAGAACGCAGC-3’) and ITS4R (5’-TCCTCCGCTTATTGATATGC-3’) with overhang adapters (forward: TCGTCGGCAGCGTCAGATGTGTATAAGAGACAGNNNNGCATCGATGAAGAACGC AGC, reverse: GTCTCGTGGGCTCGGAGATGTGTATAAGAGACAGNNNNTCCTCSSCTTATTGATA TGC) to allow the successive addition of multiplexing index-sequences. First step PCR and their thermal cycling conditions were as follows: 98°C for 3 min followed by 98°C for 30 s, 55°C for 30 s and 72°C for 30 s (25 and 30 cycles for 16S rRNA and ITS genes, respectively) and a final extension for 10 min at 72°C. The PCR products of the first step were used as a template for the second step of PCR. The second PCR amplification added multiplexing index sequences to the overhang adapters using a unique combination of primers for each sample. Thermal cycling conditions were as follows: 98°C for 3 mn followed by 98°C for 30 s, 55°C for 30 s and 72°C for 30 s (8 and 10 cycles for 16S rRNA and ITS genes, respectively) and a final extension for 10 min at 72°C. PCR products from the second step were deposited on a 2% agarose gel to validate amplification and amplicons size. Amplicon products were purified using HighPrep™ PCR Clean Up System (AC-60500, MagBio Genomics Inc., USA) paramagnetic beads using a 0.65:1 (beads:PCR product) volumetric ratio to eliminate DNA fragments below 100 bp in size and primers. Samples were normalized using SequalPrep Normalization Plate (96) Kit (Invitrogen, Maryland, MD, USA) and pooled using a 5 µl volume for each sample. The pooled sample library was concentrated using DNA Clean and Concentrator™-5 kit (Zymo Research, Irvine, CA, USA). The pooled library concentration was determined using the Quant-iT™ High-Sensitivity DNA Assay Kit (Life Technologies). The final pool concentration was adjusted to 4 nM before library denaturation and sequencing. Amplicon sequencing was performed on an Illumina MiSeq platform using Reagent Kit v2 [2 x 300 cycles] (Illumina Inc., CA, US). Demultiplexing and trimming of Illumina adapters and barcodes was done with Illumina MiSeq Reporter software (version 2.5.1.3).

The 16S rRNA gene and ITS amplicon sequences were analyzed internally using a Python notebook (available upon request). Briefly, using PEAR with default settings, forward and reverse sequences were assembled. Additional quality checks were conducted using the QIIME pipeline and short sequences were removed (< 400 bp for 16S and < 300 bp for ITS). Reference based and *de novo* chimera detection, as well as clustering into operational taxonomic units (OTUs, hereafter called taxa) were executed using VSEARCH and the adequate reference databases (Greengenes’ representative set of sequences for 16S rRNA gene and UNITE’s ITS2 reference dynamic dataset for ITS). We deliberately choose the OTU analysis over the Amplicon Sequence Variant analysis (ASVs) for consistency reasons, since the latter pipeline is currently not available for fungal ITS sequences. The identity thresholds were appropriately set at 94 % for 16S and 97 % for ITS based on our routine internal calibration controls (mock communities) in order to get the most accurate resolution. For simplicity reasons, we refer only to 16S rRNA gene amplicon sequences as “bacterial”, since archaeal sequences were extremely rare. Taxonomy was allocated using UCLUST and the latest released Greengenes database (v.05/2013). For ITS, the taxonomy assignment was executed using BLAST algorithm and the UNITE reference database (v.7-08/2016). A summary table is provided in supporting information to present the sequenced samples and the number of sequences recovered (Table S1). Sequences have been submitted to the Sequence read Archive public repository (SRA, https://www.ncbi.nlm.nih.gov/sra), under the following accession numbers for the 16S rRNA gene amplicon dataset: SUB9063594; and for the ITS2 amplicon dataset: SUB9104701

### Alpha-diversity analysis of microbial communities

To explore the sequencing completeness in terms of diversity recovery per sample, individual raw rarefaction curves were calculated using the “*vegan*” package in R (Fig. S1). After careful evaluation of the rarefaction curves, we identified a series of problematic samples coming from the same column in the 96-well plate, harboring significantly higher sequences counts but relatively lower OTU discovery rates than the others (*p* < 0.001). In order to avoid biasing our conclusions, we decided not to consider those samples for further analyses. Due to the unevenness of sequencing depth and its consequences on diversity assessment, we applied random re-sampling for normalization of samples in each data set. The samples were rarefied at 14,000 for both bacteria and fungi. Those levels are considered to be appropriated for accurate alpha-diversity estimations based on best practices guidelines (Schöler et al., 2017). Alpha-diversity analysis was perform using the following indices: observed taxa richness (S), estimated richness (Chao-1), ACE (Abundance-based Coverage Estimator), Simpson index (1-D, D = Dominance), Shannon index (H) and Equitability (J = H/ln(S)). Alpha-diversity indices were exported for bacteria and fungi. An additional PERMANOVA on the six diversity indices taken together was also performed to detect the overall effect of factors on bacterial and fungal diversity.

### Beta-diversity analysis of microbial communities

For the beta diversity analysis, we used the rarefied dataset. In order to consistently analyze both bacterial and fungal profiles with the same method, we choose the Bray-Curtis dissimilarity index to generate the distance matrices (package ‘*vegan*’, Dixon, 2003). The Bray-Curtis dissimilarity index was preferred over other metrics (e.g. UniFrac) as ITS sequences cannot be aligned to obtain a phylogenetic distance. We first assessed with PERMANOVA the significance of each factor on the structure of bacterial and fungal community. As only the breeding type and the block had a significant effect, a partial distance-based redundancy analysis (db-RDA) was used with the following model: breeding type + Condition (block) (‘*capscale*’ function, package ‘*vegan*’, 10,000 permutations).

### Network analysis

We investigated the structure of the modern and ancient wheat varieties rhizosphere microbiota using a network approach based on edge arithmetic (Jacquiod et al. 2020) to focus on OTU correlations that are specific of each type of breeding. We first calculated two separated partial correlation matrices amongst dominant rhizosphere OTUs (total summed counts = 100 minimum, occurrence = 25/51 samples), one for the modern and one for the ancient varieties using Poisson Log Normal models (PLN, package ‘*PLNmodels*‘, Chiquet et al. 2019). This method allows the combining of several datasets, thus enabling the integration of both bacterial and fungal sequencing data together using the ‘TSS’ offset criteria (Total Sum Scaling). The following model was built to account for the block effect for both modern and ancient varieties “∼ 1 + block + offset”. The PLN models were validated by using the Bayesian Information Criterion (BIC) to determine the most appropriated sparsifying penalty levels (BIC, R^2^ ancient = 0.98; R^2^ modern = 0.97). We then applied edge arithmetic with the ‘*igraph’* package functions (Csardi and Nepusz, 2006). Briefly, we intersected both modern and ancient networks to systematically determine whether correlations were common to both breeding types or specific of one breeding type. For visualization, the two networks were then merged and opened in the ‘*Cytoscape*’ software (Shannon et al. 2003). The complexity of networks was investigated by means of the degree index, the node betweenness and the edge betweenness.

### Mycorrhizal colonization

Two root systems per plot were collected and pooled together 29 weeks after sowing, from the 22^th^ to the 24^th^ of May 2017. Root material was washed thoroughly and prepared for staining as described in Vierheilig et al. (1998). Roots were cleared at 90°C for 5 to 10 min in 10% KOH, placed in black ink (5% in acetic acid) for 5 min at 90°C for coloration, rinsed with water and put for 25 min in 8% acetic acid at room temperature. Roots were rinsed again with water, covered with pure glycerol and stored at 4°C. For microscopic observation, roots were cut into 1 cm fragments and 15 fragments were placed in glycerol between slide and slip cover four times per sample. There were two batches of microscope slides for convenience of counting, called “Series” in the statistical analysis. Mycorrhization rates were assessed according to Trouvelot et al. (1986) and expressed as mycorrhizal colonization i.e. frequency of root fragments with mycorrhizal structures at the root system scale (F), intensity of the mycorrhizal colonization at the root system scale (M) or restricted to myccorhizal root fragments (m), arbuscule abundance at the root system scale (A) or restricted to mycorrhizal root fragments (a).

#### Univariate statistical analysis

The statistical analysis of univariate data, including alpha-diversity indices and mycorrhiza traits, was performed in Rstudio software (RStudio Team, 2020). Normality and homoscedasticity of the data was assessed using Shapiro and Bartlett test respectively using R default functions. Normally distributed data were analyzed with construction of an ANOVA model, significance was assessed using the D’Agostino test of skewedness on the residual variance (package ‘*moments*’, Komsta and Novomestky, 2015), followed by a post-hoc Tukey’Honest Significant Detection test (Tukey’s HSD, p<0.05, package ‘*agricolae*’, De Mendiburu, 2017). Non-parametric data were analyzed with a Kruskal-Wallis test followed by a post-hoc False Discovery Rate test correction to account for multiple testing (FDR, *p* < 0.05, package ‘*agricolae*’). We present the percentage of variance explained and associated *p*-values from the ANOVA models if variables are normally distributed, and only the *p*-value from the Kruskal-Wallis test if not normally distributed.

## RESULTS

### Effects of breeding types and inputs on bacterial community

Breeding type (ancient *vs* modern varieties) had a significant effect on all bacterial alpha diversity indices, including the richness (explaining 15.56% of the total variance, *p* < 0.001), Chao-1 (6.9%, *p* = 0.025), ACE (5.9%, *p* = 0.044), Simpson reciprocal (*p* = 0.044), Shannon (*p* = 0.002) and Equitability (*p* = 0.012) (Tables 1, Table S2). The presence/absence of inputs had a barely significant effect on community evenness, including the Simpson reciprocal (*p* = 0.048) and Equitability (*p* = 0.057) (Tables 1 and S2). The interaction between breeding type and inputs had a significant effect on richness (7.7%, *p* = 0.011), ACE (6.8%, *p* = 0.031), Simpson reciprocal (*p* = 0.013), Shannon (*p* = 0.001) and Equitability (*p* = 0.006). The different varieties inside each breeding type was the main source of variation for richness (46.1%, *p* = 0.009), Chao1 (49.9%, *p* = 0.015) and ACE (47.5%, *p* = 0.028) indices. When the six diversity indices were analyzed simultaneously in a PERMANOVA (Table 2), we found an overall significant effect of the breeding type (7.4%, *p* = 0.013) and its interaction with chemical inputs (5.7, *p* = 0.034). The variety error term inside breeding type was also significant (48.8%, *p* = 0.015). The overall multivariate model explained 63.5% of the variance.

**Table 1:** Analysis of variance and Kruskal-Wallis tests for bacterial alpha diversity indices. F and chi-square (χ²) values are given, with asterisks indicating the significance of effects. “Block” is the factor identifying the three repeated blocks in the experimental design. “Inputs:Breeding type:Variety” is an error term for the variance explained by the variety inside each breeding type. For non-parametric data, Kruskal-Wallis tests were done individually on each factor or factor combinations. The directions of significant effects are indicated. w, presence of inputs; w/o, absence of inputs; Mod., modern varieties; Anc., ancient varieties; ᐧ, *p* < 0.10; *, *p* < 0.05; **, *p* < 0.01; ***, *p* < 0.001.

| Factors | Richness (F) | Chao-1 (F) | ACE (F) | Simpson reciprocal ($\chi^2$) | Shannon ($\chi^2$) | Equitability ($\chi^2$) |
| --- | --- | --- | --- | --- | --- | --- |
| Inputs | 0.063 | 1.688 | 0.207 | 3.914* | 1.278 | 3.622 . |
| Breeding type | 14.81*** | 5.586* | 4.436* | 4.077* | 9.578** | 6.300* |
| Block | 0.043 | 0.444 | 0.437 | 5.233 . | 2.611 | 4.972 . |
| Inputs x Breeding type | 7.363* | 3.412 . | 5.151* | 10.83* | 15.64** | 12.50** |
| Inputs x Breeding type/Variety | 2.740** | 2.528* | 2.247* | 10.10 | 14.31 | 11.64 |
| Direction of effects | Mod. > Anc. | Mod. > Anc.<br>w/o > w in Anc. |  | w/o > w<br>Mod. > Anc.<br>w/o > w in Anc. | Mod. > Anc.<br>w/o > w in Anc. | w/o > w in Anc. |
| R<sup>2</sup> (variance explained) | 0.695 | 0.642 | 0.616 | na | na | na |

**Table 2:** Permutational analysis of variance for bacterial and fungal alpha diversity based on the six diversity indices (Richness, Chao-1, ACE, Simpson reciprocal, Shannon and Equitability) and for parameters describing mycorrizal colonization (F, M, m, A and a, see Materials and Methods for a description). F values are given, with asterisks indicating the significance of effects. ᐧ, *p* < 0.10; *, *p* < 0.05; **, *p* < 0.01; ***, *p* < 0.001.

| PERMANOVA | R<sup>2</sup> (Bacteria) | R<sup>2</sup> (Fungi) | R<sup>2</sup> (Myco) |
| --- | --- | --- | --- |
| Input | 0.014 | 0.004 | 0.008 |
| Breeding | 0.074* | 1.7E-05 | 0.028** |
| Input x Breeding | 0.057* | 0.105* | 0.011 |
| Input x Breeding/variety | 0.478* | 0.118 | 0.188** |
| Block | 0.012 | 0.046 | 0.028* |
| Residual | 0.365 | 0.727 | 0.737 |
| R<sup>2</sup> (variance explained) | 0.635 | 0.273 | 0.263 |

Regarding the direction of the effects of factors on bacterial diversity, we found that, considering both treatments with and without inputs together, modern varieties had richer and more even communities as compared to ancient varieties (Fig. 1). Considering both ancient and modern varieties together, the presence of inputs had significant effects only on evenness indices, with a slight increase in dominance of some taxa. However, the addition of inputs was responsible for a strong and significant decrease in diversity for ancient varieties for richness, Simpson reciprocal, Shannon and Equitability, but it had no effect on modern varieties (Fig. 1).

**Figure 1:**
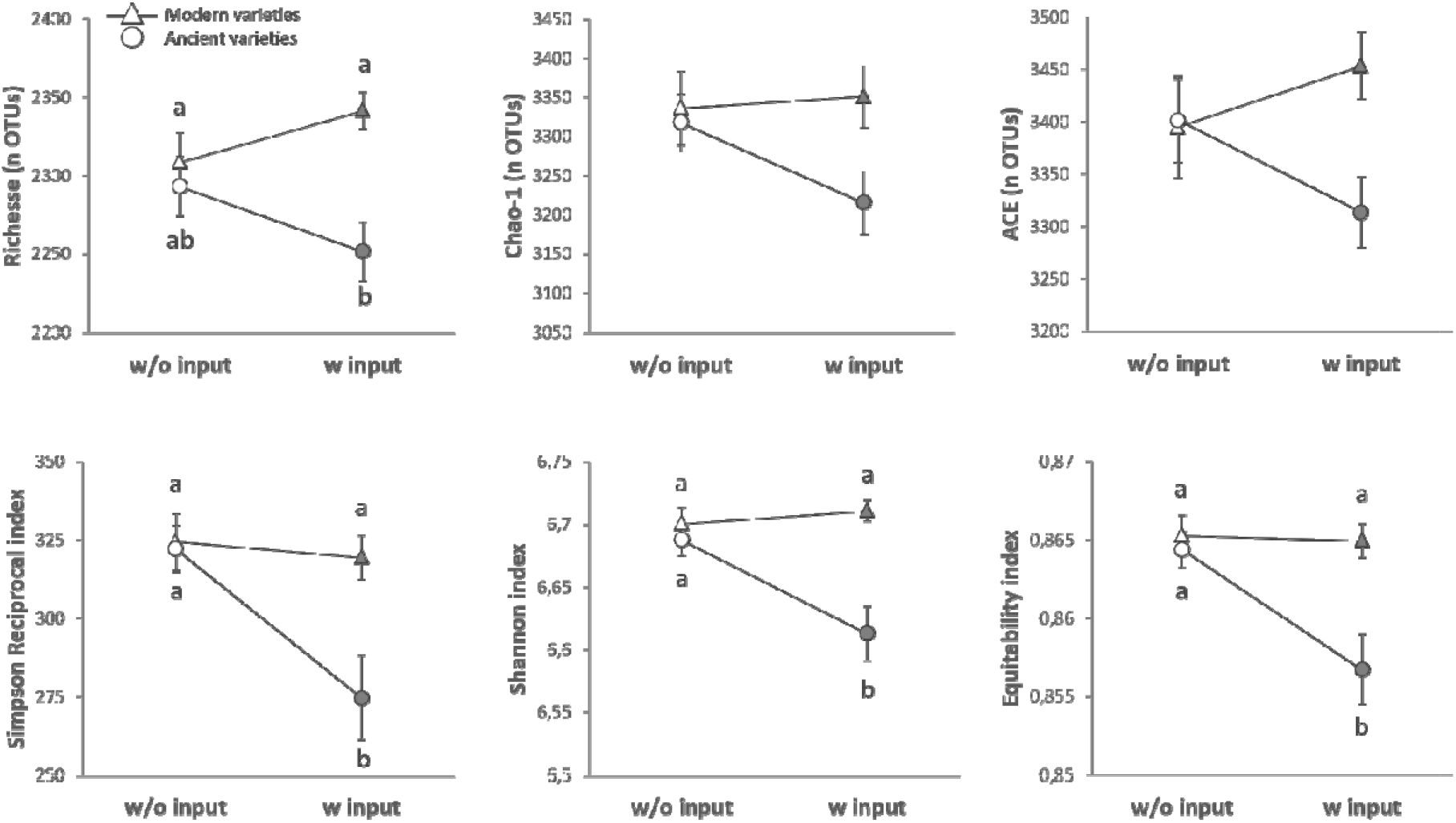
Alpha-diversity of the rhizosphere bacterial community (16S rRNA gene amplicon sequencing), presented as reaction norms of ancient and modern wheat genotypes to the environment modification by inputs. Panel are respectively showing: the richness (A), the Chao-1 estimator (B), the ACE estimator (C), the Simpson reciprocal index (D), the Shannon index (E) and the Equitability of Pielou (F). Data is shown as the interaction between the breeding type and the presence/absence of inputs (w/o: without inputs; w: with inputs). Significance was inferred with an ANOVA under the Tukey’s HSD post-hoc test for normally distributed data (Honest Significant Detection, *p* < 0.05). Non-parametric data were analyzed with a Kruskal-Wallis test under False Discovery Rate post-hoc correction (FDR, *p* < 0.05).

The PERMANOVA on bacterial community beta diversity (Table 3) showed that the breeding type (2.7%, *p* = 0.036) and the block (6.5%, *p* < 0.001) but not the inputs had a significant effect (1.8%, *p* = 0.701). From this result, we profiled the structural changes in bacterial communities using a distance-based partial redundancy analysis on Bray-Curtis dissimilarities for breeding type only (the block effect was added as an error term). The total variance explained was 2.3% (*p =* 0.016) and the first constrained component (CAP1) and first non-constrained component (MDS1) were explaining 2.5 and 4.2% of variance, respectively (Fig. 2A). Ordination plots indicated that bacterial community structure differed between breeding types.

**Figure 2:**
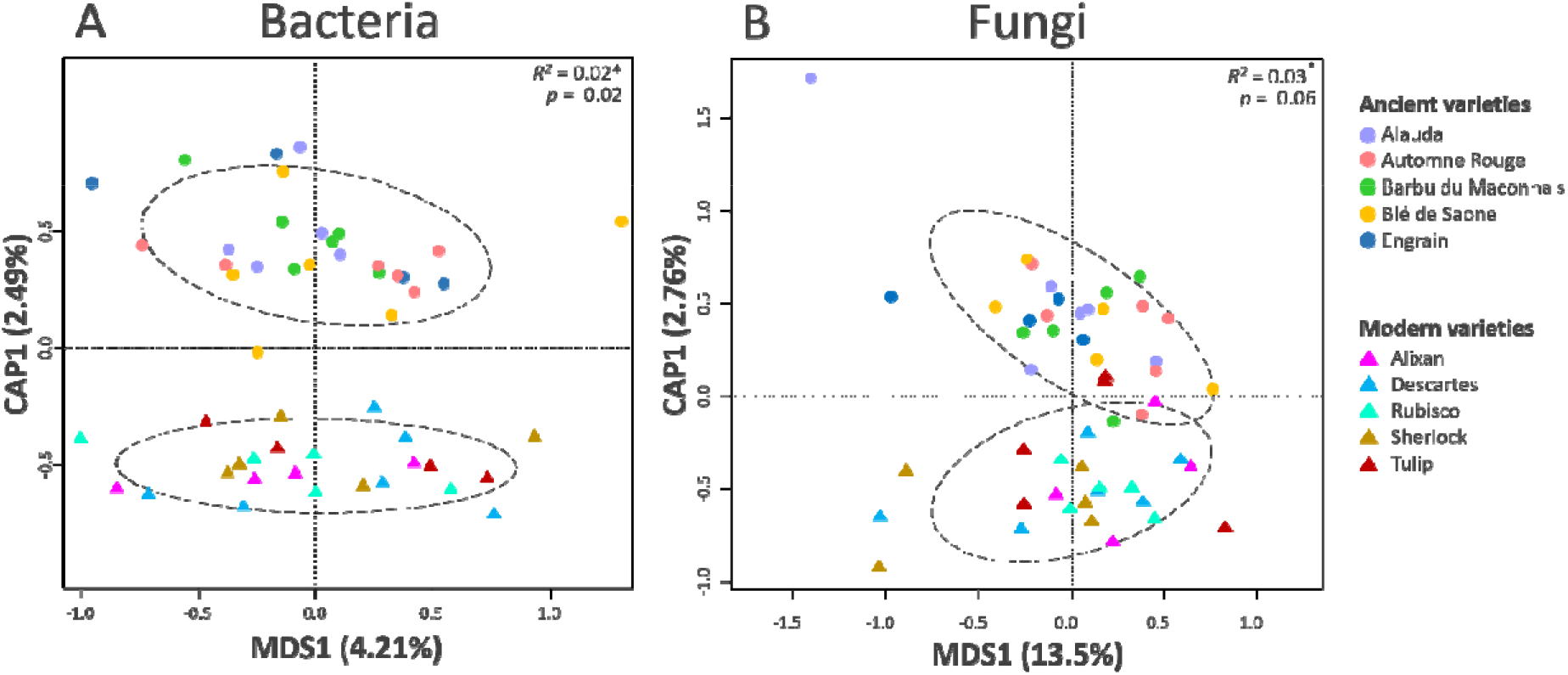
Beta-diversity analysis of the bacterial (A) and fungal (B) communities using partial distance-based redundancy analysis (db-RDA). Partial db-RDA were used to output the following constrained models: breeding type + Condition (block) (10,000 permutations).

**Table 3:** Permutational analysis of variance for bacterial (16S) and fungal (ITS) beta diversity. F values are given, with asterisks indicating the significance of effects. The directions of effects are indicated. Mod., modern varieties; Anc., ancient varieties; ᐧ, *p* < 0.10; *, *p* < 0.05; **, *p* < 0.01; ***, *p* < 0.001.

| Factors | Bacteria (16S rRNA gene) | Fungi (ITS2 fragment) |
| --- | --- | --- |
| Inputs | 0.018 | 0.022 |
| Breeding type | 0.027* | 0.025* |
| Block | 0.065*** | 0.054* |
| Inputs x Breeding type | 0.019 | 0.017 |
| Inputs x Breeding type/Variety | 0.308 | 0.347* |
| Residual | 0.564 | 0.535 |

### Effects of breeding types and inputs on fungal community

Breeding type had no significant effect on fungal diversity (Table 4, Table S3). The presence/absence of inputs showed a significant effect on the Shannon diversity (6.7%, *p* = 0.044). The interaction between breeding type and inputs had a significant effect on the Chao-1 estimation (13.9%, *p* = 0.025). The effect of the different varieties inside each breeding type had no significant effect, despite a relatively high percentage of variance (Table S3). When the six diversity indices were analyzed simultaneously in a PERMANOVA (Table 2), we found no significant effects of inputs (0.4%, *p* = 0.765), breeding type (0.02%, *p* = 0.995), variety (11.8%, *p* = 1.000) and block (4.6%, *p* = 0.444) alone, but the breeding x inputs interaction was responsible for an overall effect on fungal diversity indices, with 10.5% of explained variance (*p* = 0.053). The overall multivariate model explained 27.3% of the variance.

**Table 4:** Analysis of variance and Kruskal-Wallis tests for fungal alpha diversity indices. F and chi-square (χ²) values are given, with asterisks indicating the significance of effects. “Block” is the factor identifying the three repeated blocks in the experimental design. “Inputs:Breeding type:Variety” is an error term for the variance explained by the variety inside each breeding type. For non-parametric data, Kruskal-Wallis tests were done individually on each factor or factor combinations. The directions of significant effects are indicated. w, presence of inputs; w/o, absence of inputs; Mod., modern varieties; Anc., ancient varieties; ᐧ, *p* < 0.10; *, *p* < 0.05; **, *p* < 0.01; ***, *p* < 0.001.

| Factors | Richness (F) | Chao-1 (F) | ACE (F) | Simpson reciprocal ($\chi^2$) | Shannon (F) | Equitability (F) |
| --- | --- | --- | --- | --- | --- | --- |
| Inputs | 1.002 | 0.005 | 0.048 | 0.746 | 4.457* | 3.601 . |
| Breeding type | 0.001 | 0.012 | 0.004 | 1.141 | 0.899 | 1.124 |
| Block | 0.924 | 1.148 | 0.677 | 4.557 . | 6.571** | 5.987** |
| Inputs x Breeding type | 2.845 | 5.624* | 3.097 . | 2.590 | 3.035 . | 1.413 |
| Inputs x Breeding type/Variety | 0.298 | 0.282 | 0.282 | 10.40 | 1.049 | 1.277 |
| Direction of effects |  | w/o < w in Mod.<br>w/o > w in Anc. | w/o < w in Mod.<br>w/o > w in Anc. |  | w/o > w<br>Mod. < Anc. in w/o | w/o>w |
| R<sup>2</sup> (explained variance) | 0.272 | 0.308 | 0.243 | na | 0.578 | 0.579 |

Regarding the direction of effects, we found no effect of the breeding type or of the presence of inputs alone (Fig. 3). Ancient varieties in the absence of inputs had the highest observed fungal diversity (not significant after post-hoc correction for multiple comparison, likely due to the interference of the block effect). For most of the diversity indices, the addition of inputs induced a decrease in fungal diversity for ancient varieties and, conversely, an increase in diversity in modern varieties (Fig. 3). The significant interaction was illustrated by the crossing of all reaction norms, which was only supported statistically for the Chao-1 index.

**Figure 3:**
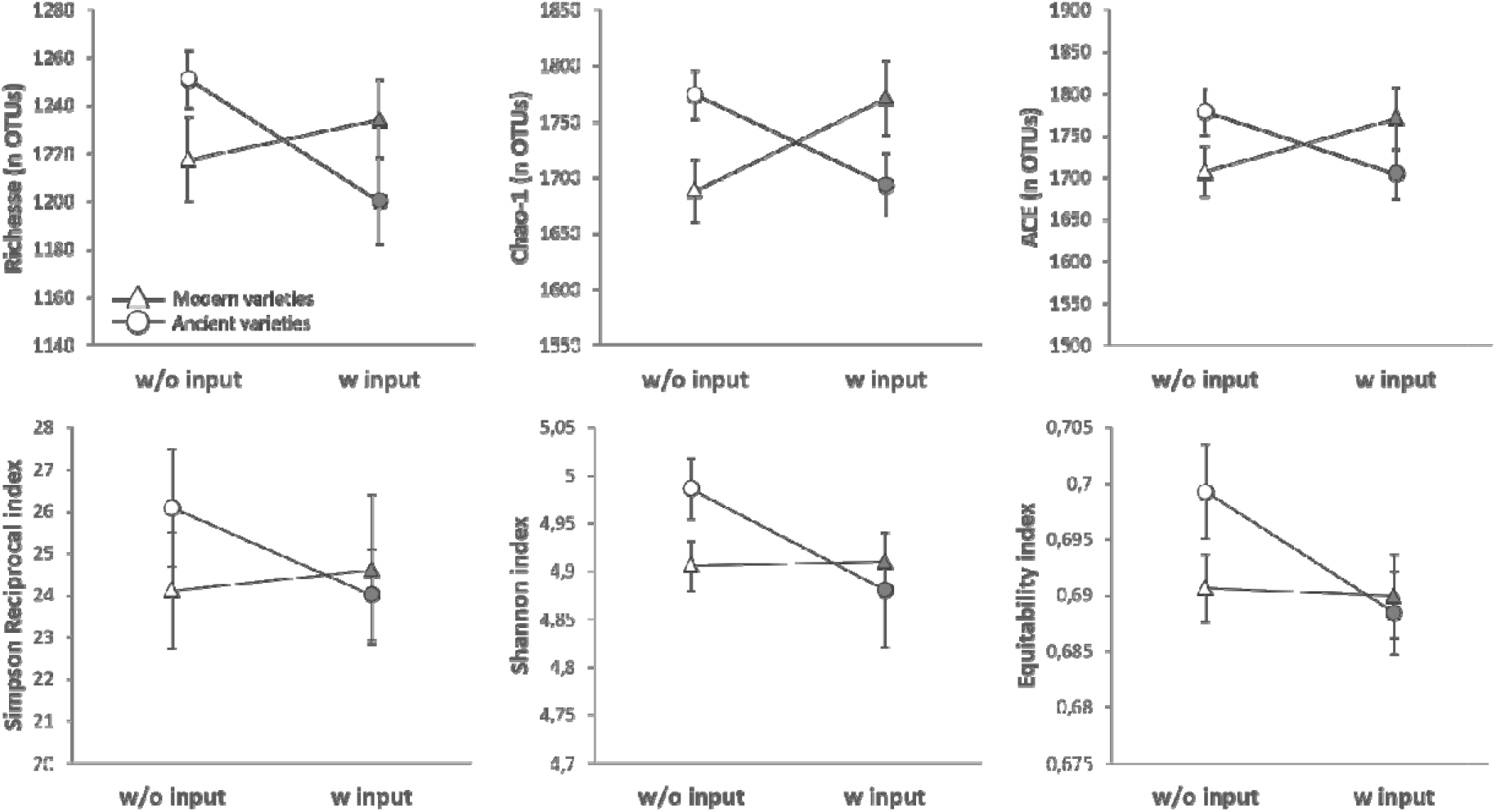
Alpha-diversity of the rhizosphere fungal community (ITS2 gene amplicon sequencing), presented as reaction norms of ancient and modern wheat genotypes to the environment modification by inputs. Panel are respectively showing: the richness (A), the Chao-1 estimator (B), the ACE estimator (C), the Simpson reciprocal index (D), the Shannon index (E) and the Equitability of Pielou (F). Data is shown as the interaction between the breeding type and the presence/absence of inputs (w/o: without inputs; w: with inputs). Significance was inferred with an ANOVA under the Tukey’s HSD post-hoc test for normally distributed data (Honest Significant Detection, *p* < 0.05). Non-parametric data were analyzed with a Kruskal-Wallis test under False Discovery Rate post-hoc correction (FDR, *p* < 0.05).

The PERMANOVA on fungal community beta diversity showed that the breeding type (2.5%, *p* = 0.052), the block (5.4%, *p* =0.011) and variety (34.7%, *p* = 0.020) had significant effects, but not the inputs (2.2%, *p* = 0.14). Based on this result, we profiled the structural changes in fungal communities using a distance-based partial redundancy analysis based on Bray-Curtis dissimilarities for breeding types alone (the block effect was added as an error term). The total variance explained was 3.0% (*p =* 0.057), and the first constrained component (CAP1) and first non-constrained component (MDS1) were explaining 2.76 and 13.50% of variance, respectively (Fig. 2B). As for bacterial community structure, ordination plots showed a differentiation trend in the fungal community structure between modern and ancient varieties while inputs addition had no effect.

### Network analysis

The PLN models successfully converged into stable networks for the modern varieties (BIC R^2^ = 0.97) and ancient varieties (BIC R^2^ = 0.98). The modern network featured 178 OTUs (nodes) and 452 edges (correlations) of which 245 were positive and 207 negative. The ancient network featured 185 OTUs (nodes) and 402 edges (correlations) of which 238 were positive and 164 negative. The comparison between the two networks revealed a similar level of node degree (Fig. 4A), but a significantly higher level of centrality in the ancient network based on average betweenness (either on nodes: *p* = 0.05; and on edges *p* = 4.93.10^-14^; Fig. 4BC). The positive-to-negative edge ratio was in favour of more positive correlations in the ancient network (1.40 vs 1.14). When combining both networks and highlighting the position of common and unique edges found either in modern or ancient varieties, a clear distinction was found, with a clearly different interaction structure depending on the breeding type (Fig. 4A). In total, this combined network featured 252 unique OTUs, amongst which 108 were common to both breeding type, while 76 were unique of the ancient varieties and 68 of the modern varieties. Most OTUs were affiliated to Ascomycota (55%), which clearly dominated both networks.

**Figure 4:**
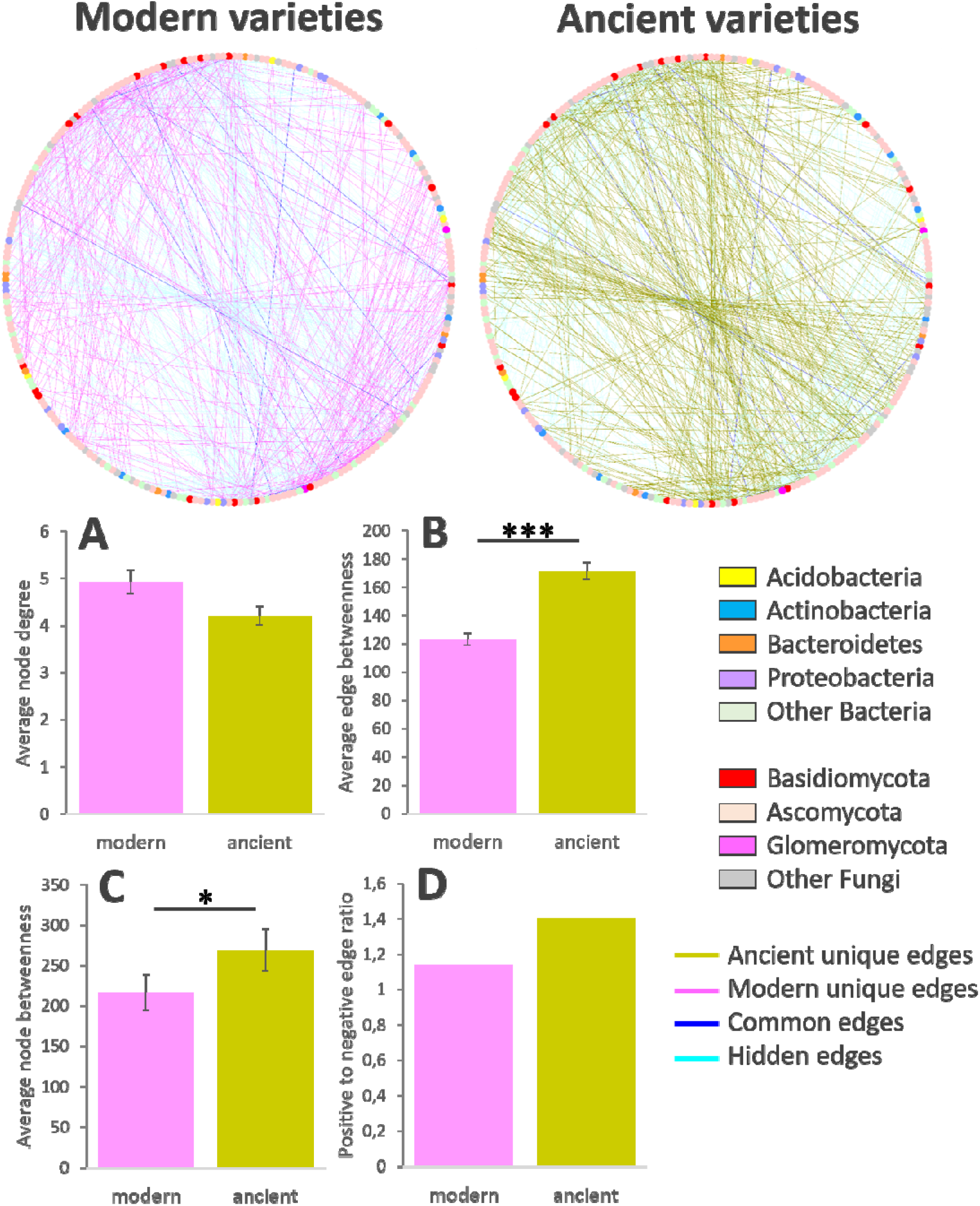
Network analysis of the rhizosphere microbiota associated to the modern and ancient wheat varieties. The networks were constructed using dominant OTUs (sum min = 100 counts, occurrence = 25/51 for bacteria, and 25/50 for fungi) and based on partial correlations obtained with Poisson Log Normal models combining the bacterial and fungal datasets together (∼1 + block + offset). For visualization, both networks were merged into one, and shown twice with edges that are unique of modern varieties highlighted on the left network (pink), and edges that are unique of the ancient varieties highlighted on the right network (khaki). Common link are highlighted in both networks in deep blue. Hidden links are corresponding to modern/ancient edges respectively not shown in each network. Barcharts represent average network complexity indices estimated from edges or nodes (Kruskal-Wallis test, *p* < 0.05).

### Mycorrhizal colonization response to breeding types and inputs

The frequency of mycorhization, F, (number of mycorrhizal root fragments divided by the total number of observed fragments) was affected neither by the breeding type, the inputs nor their interaction (Table 5 and S4). Breeding types had an impact on the intensity of mycorrhizal colonization in root system M (explaining 4.3% of total variance, *p =* 0.005), in mycorrhizal root fragments m (4.5%, *p =* 0.009), as well as on arbuscule abundance in root system A (*p =* 0.018) and in mycorrhizal root fragments a (5.6%, *p=* 0.002). Inputs had an impact only on the intensity of mycorrhizal colonization in root system M (explaining 2.6% of the total variance, *p=* 0.027) and in mycorrhizal root fragments m (2.6%, *p=* 0.046). The interaction between breeding type and inputs was only significant on arbuscular abundance in root system A (*p* = 0.032, Table 5). When performing a PERMANOVA on all mycorrhiza indices, a significant breeding effect was detected, in favor of higher index values for the modern varieties (2.8%, *p* = 0.008, Table 2). The overall multivariate model explained % of the variance. Regarding the direction of effects, mycorrhiza colonization was higher in modern varieties compared to ancient ones (Fig. 5ABCD) and the application of inputs decreased the intensity of mycorrhizal colonization in the root system and root fragment (Fig. 5EF).

**Table 5:**
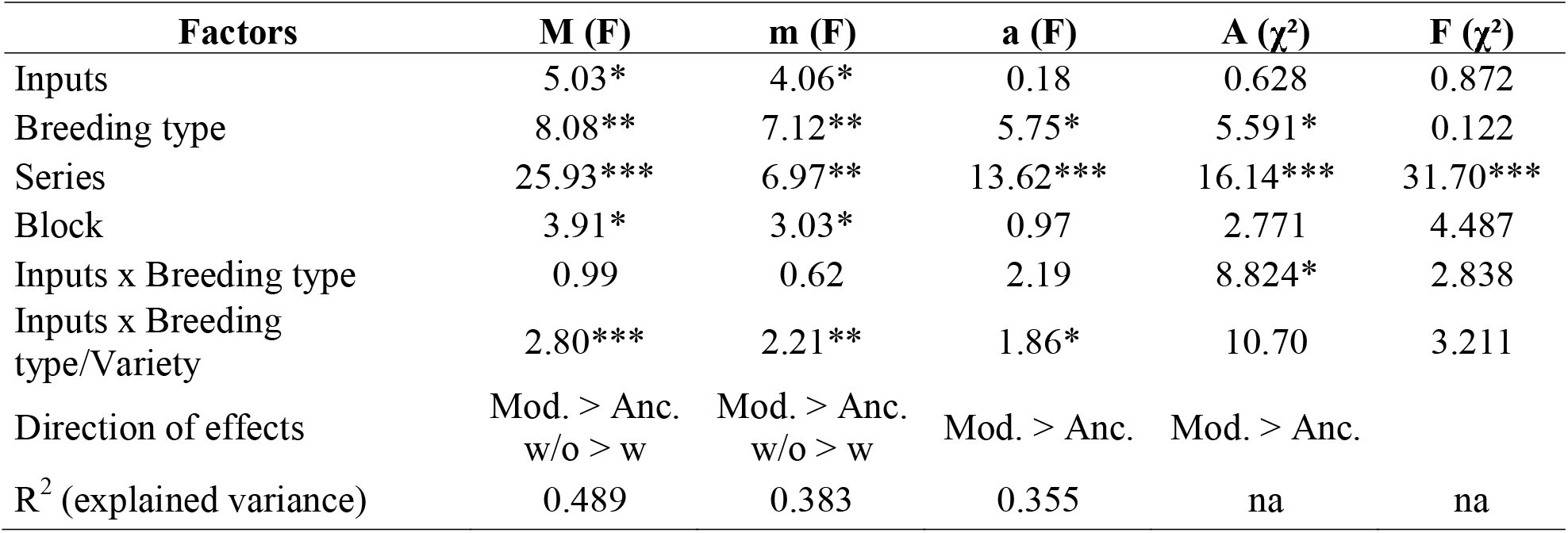
Analysis of variance and Kruskal-Wallis tests for myccorhiza indices. F and chi-square (χ²) values are given, with asterisks indicating the significance of effects. “Block” is the factor identifying the three repeated blocks in the experimental design. “Series” refers to the two different batches of staining of mycorrhiza. “Inputs:Breeding type:Variety” is an error term for the variance explained by the variety inside each breeding type. For non-parametric data, Kruskal-Wallis tests were done individually on each factor or factor combinations. The directions of significant effects are indicated. w, presence of inputs; w/o, absence of inputs; M, modern varieties; A, ancient varieties; ᐧ, *p* < 0.10; *, *p* < 0.05; **, *p* < 0.01; ***, *p* < 0.001.

## DISCUSSION

Modern varieties of wheat were selected and generally grown with synthetic chemical inputs whereas ancient varieties were selected and grown without this kind of inputs, in organic farming systems. This correlation between variation in genotype and environment prevent to assess the relative importance of individual factors (E and G) and their interaction (G×E) in plant-microbiota relationships. In addition, comparison between modern and ancient varieties are generally made in controlled conditions in the absence of inputs or in nutrient depleted soils, which prevents any realistic conclusion on the evolution of plant-microbiota relationships in the field, since modern varieties are grown with inputs. Our results are a first attempt to quantify these environmental and genotypic effects independently, in the field.

### Effects of inputs on plant-microbiota interactions

The presence/absence of inputs – here considered as our environmental conditions “E” – had a low impact on plant-microbiota relationships. When considered independently of the breeding type, inputs had only a slight significant effect on microbial alpha-diversity, via a lowering of specific evenness indices when applied, thus indicating an increased dominance of some microbial OTUs (Fig. 1 & 3; Tables 2 & 4; Tables S2 & S4). The intensity of mycorrhizal colonization was higher in the absence than in the presence of inputs (M and m, Table 5 and Fig. 5). However, this had no consequence on the intensity of arbuscule development (A and a, Table 5 and Fig. 5), and thus likely no functional impact. The presence/absence of inputs also had no effect on the overall composition of fungal and bacterial communities in the rhizosphere (Fig. 2). Taken together, these results support our first hypothesis that the presence of inputs decreases the microbial diversity in the rhizosphere. This decrease in diversity in the presence of inputs was expected for several reasons. First, the addition of N fertilizer can induce a shift in bacterial communities. In line with our results, Grunert et al. (2019) reported that the addition of an inorganic fertilizer (struvite) reduced the diversity (Pielou equitability, Shannon and Simpson reciprocal) of tomato rhizosphere microbiota. In a study with two sites where N was added for 27 and 8 years, the authors found that bacterial community structure was highly responsive to N additions, but the diversity of bacterial community did not have a consistent response (Ramirez et al., 2010). As a matter of fact, the shift in bacterial community composition is mainly observed in soils with low C and N concentrations (Ramirez et al., 2012). The C and N concentrations in our soil presented a medium value, which could explain the weak effect of nitrogen fertilization effect. Second, the effect of fungicide could have suppressed some fungal taxa. It has been shown that commonly used fungicides with foliar application had moderate but significant effect on the composition of fungal communities in the wheat phyllosphere (Karlsson et al., 2014). While fungicides are supposed to target specific fungal pathogens, the impact on fungal communities has been already observed (Esmaeili Taheri et al., 2015). Third, the effect of herbicide could have modified the weed community, which in turn could have influenced microbial community, although this mechanism has not been studied yet, to our knowledge. Our weak observations can also be explained by the fact that inputs are often reported to affect *bulk soil* microbial communities, whereas *rhizosphere* microbial communities are more dependent on plant factors than on soil properties (Grunert et al., 2019).

**Figure 5:**
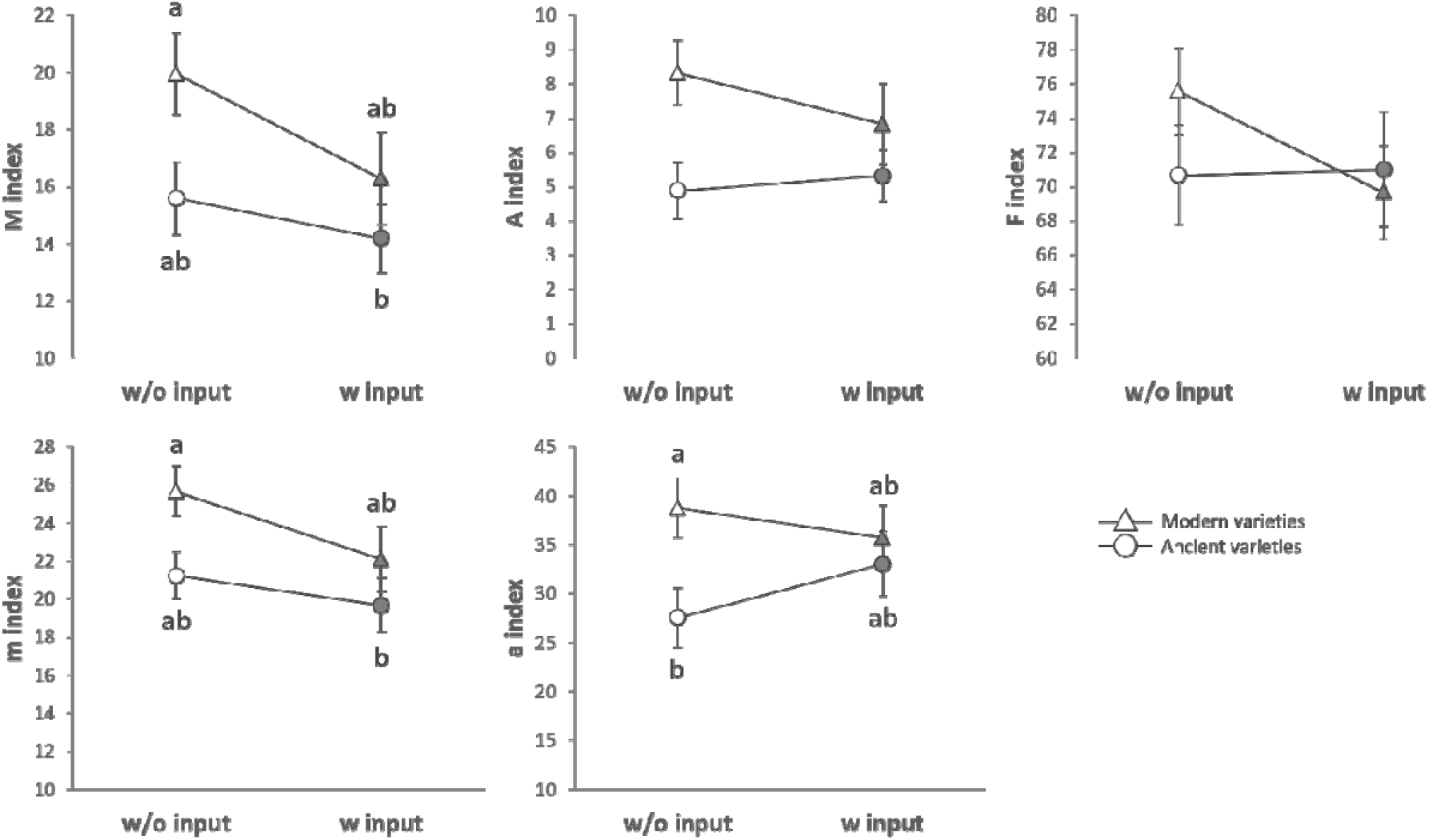
Mycorrhizal colonization for breeding types and inputs. Panel are respectively showing: the intensity of mycorrhizal colonization in root system (A), the intensity of mycorrhizal colonization in root fragment (B), the arbuscular abundance in root system (C), the arbuscular abundance in root fragment (D) in response to the breeding types (whatever the presence of inputs) and the intensity of mycorrhizal colonization in root system (E) and the intensity of mycorrhizal colonization in root fragment (F) in response to inputs (whatever the breeding type). Significance was inferred with an ANOVA under the Tukey’s HSD post-hoc test for normally distributed data (Honest Significant Detection, *p* < 0.05). Non-parametric data were analyzed with a Kruskal-Wallis test under False Discovery Rate post-hoc correction (FDR, *p* < 0.05).

### Effects of breeding on plant-microbiota interactions

The breeding type – here considered as the genotype G – had a strong effect on rhizosphere microbial communities, which was dependent on the microbial taxa (bacteria or fungi), meaning that plant-microbiota relationships were differently influenced by breeding from ancient to modern varieties. When considering the impact of the genotype independently of the use of inputs, we found that breeding type affected bacterial diversity, with a percentage of explained variance from 5.9 to 15.6% of the variance according to the index (Table S2) and 7.4% of the variance when all indices were analyzed together (Table 2). The variety inside each breeding type was the strongest determinant of bacterial diversity, explaining 46.1 to 49.9% of the total variance, and 48.8% when all indices were analyzed together, stressing the importance of genotype. Conversely, the breeding type had no effect on fungal diversity (Table S3). However, all mycorrhiza indices (except F) were higher with modern varieties compared to ancient ones (Table 5, Table 2, Fig. 5). The breeding type also affected slightly the overall composition of fungal and bacterial communities (Table 3, Fig. 2), but its strongest effect was noticed on the structure of the rhizosphere microbiota network (Fig. 4). Indeed, the different breeding types featured completely distinct co-occurrence links amongst the dominant microbial OTUs, which resulted in a significantly less complex network for the modern varieties compared to the ancient ones. Therefore, our results partially invalidated the second hypothesis that modern genotypes have lost their ability to establish interactions with microbial species, as we evidenced that this interaction still exists and may even be reinforced in the case of mycorrhiza. However, the interaction with bacteria and other fungal members has been completely restructured into a simplified form.

Although this hypothesis is supported by several studies in the literature, there is no consensus. Among supporting results, Valente et al. (2020) found that ancient wheat varieties were more capable of interacting with beneficial plant growth rhizobacteria than modern varieties. Moreover, a decrease in the diversity of symbiotic rhizobia associations has been observed in domesticated legumes compared to wild relatives (Kim et al. 2014; Sangabriel-Conde et al. 2015). Mutch and Young (2004) reported that the ability to interact with symbionts was limited for modern pea and broad bean as compared to wild relatives of the *Vicia* and *Lathyrus* genera in a non-agricultural soil without inputs. Other studies found more nuanced results: Leff et al. (2016) reported that neither root nor rhizosphere bacterial communities were affected by sunflower domestication, but domestication did affect the composition of rhizosphere fungal communities. Brisson et al. (2019) reported similar Shannon index for prokaryotic or fungal communities in teosinte, inbred and modern varieties; they observed that co-occurrence network of microbiota of inbred maize lines’ rhizosphere were significantly closer from those of the teosintes than to the modern hybrids. Opposed results also exist: Kinnunen-Grubb et al. (2020) reported that bacterial colonization in the roots of modern cultivars of wheat is faster than in ancestors. This could be explained by the fact that modern varieties could have lost their ability to specifically select for beneficial microbial partners compared to ancient varieties (Kiers and Denison, 2008).

Indeed, with six soybean cultivars representing 60 years of breeding, Kiers et al. (2007) showed that ancient varieties were more performant than modern varieties in maintaining a high fitness (seed production) when infected with a mixture of effective and ineffective rhizobia strains. This lower selectivity of modern varieties could explain the associated higher diversity in rhizosphere microbiota that we observed in the presence of inputs. However, a metaLanalysis on the effects of breeding on mycorrhizal responsiveness found that varieties released after 1950 were more mycorrhizalLresponsive (in terms of increased biomass production) than old varieties (1900-1950) and ancestors (before 1900) (Lehmann et al., 2012). So the fact that modern varieties establish either adaptive or non-adaptive interactions with soil microbiota remains an open question (see Ghalambor et al., 2007 for in-depth discussion). The aim of our experiment was not to evaluate the fitness gain due to plant-microbiota interactions, but it would be an interesting perspective.

### Effects of phenotypic plasticity on plant-microbiota interactions

We speculate that these contradictory results regarding the effect of breeding could be due to the interaction between plant genotype and the environment. Our experimental design allowed to assess whether variations in plant-microbiota interaction are affected by the interaction between the breeding type and the presence of inputs, called phenotypic plasticity. Namely, phenotypic plasticity describe the capability of a genotype to produce different phenotypes in response to variation in environmental conditions (Gause, 1947; Bradshaw, 1965). It is an ubiquitous aspect of organisms. The profile of phenotypes produced by a genotype across environments is the “norm of reaction” (Schmalhausen, 1949; Stearns, 1989).

Indeed, phenotypic plasticity – here considered as the G×E interaction – had an important impact on plant-microbiota relationships. It affected several indices of bacterial and fungal diversity (Tables 1 and 3, Tables S2 and S3, Figs. 1 and 3). For the bacterial community, this effect of phenotypic plasticity (5.7%) was in the same range as the effect of the breeding type (7.4%, Table 2). For the fungal community, the effect of the breeding type was not significant, whereas the effect of phenotypic plasticity was significant and relatively important when considering all indices together (10.5%, Table 2). However, phenotypic plasticity had no impact on the intensity of mycorrhizal colonization (Table 5) or on the overall structure of bacterial and fungal communities (Table 3). This G×E interaction was key in understanding the variations of diversity: ancient varieties had a decreased bacterial and fungal diversity in the presence compared to the absence of inputs, whereas modern varieties kept diverse rhizosphere microbial community in the presence of inputs. This contradicts our third hypothesis that the lower plant-microbiota association with modern genotypes is amplified in the presence of inputs. The literature reports many cases of interaction between genotype and environment for other plant traits and considering plasticity as a trait among others allows finding out some genetic determinant of plasticity. For example, grain yield variation in maize was found to result from the environment (a large geographic and climatic transect of North American) for 43%, genotype for 7% and G×E for 6% (see Gage et al. 2017, Supp. Fig. 4). The authors found that genomic regions that have experienced changes in allele frequency due to selection for productivity in temperate conditions explain less G×E variation than regions in which allele frequency was unaffected by selection. Loci associated with G×E interaction were mainly located in the regulatory regions of the genome (Gage et al., 2017). Regarding plant-microbiota interactions, it has been observed in *Boechera stricta* (Brassicaceae), a perennial wild mustard grown in various sites in North America, that the G×E effect on microbial community was stronger for the phyllosphere than the rhizosphere, with an impact on the Shannon index greater than the impact of the genotype itself (but the opposite was true for Chao1) (Wagner et al., 2016). We found one experiment considering both wild relatives and cultivated varieties and the manipulation of inputs as an environmental variation (Shi et al. 2019). Using one single ancient and modern varieties of soybean and rice, they reported that the rhizosphere microbiota of wild relatives are more affected by a fungicide than the cultivated ones and that the G×E interaction had a significant effect on fungal community. However, they did not discuss their results in terms of phenotypic plasticity.

## CONCLUSION

The evolution of the phenotypic plasticity of plant-microbiota interactions has been neglected up to now by microbial ecologists and geneticists. This knowledge gap makes difficult any robust conclusion on the fact that modern varieties have lost or improved their ability to interact with soil microbiota. Our results suggest that modern varieties are less sensitive to the negative effect of inputs on plant-microbiota interactions than ancient varieties. This could be explained by a loss of ability to regulate plant-microbiota interactions according to soil fertility in modern varieties, in accordance the results of Gage et al. (2017) showing that G×E variation is disproportionately controlled by regulatory mechanisms as compared with other traits. This is coherent with the fact that modern varieties were not exposed to huge variations of soil fertility such as ancient ones, as they have been selected and grown in the presence of inputs. So there were less selective constraints on the modern regarding the ability to fit with the heterogeneity of soil fertility. However, environmental pollution by synthetic chemical inputs will likely lead to grow crops with less inputs, so in environments with more heterogeneous levels of fertility. Integrating phenotypic plasticity in plant-microbiota interactions as a target in future breeding programs could be very useful to obtain varieties with high yield, but able to finely adapt their interactions with microbiota to variations in soil fertility.

## ACKNOWLEDGEMENTS

We thank AgroSup Dijon for experimental facilities. We are grateful to Jean-Philippe Guillemin, Wilfried Queyrel and Etienne Gaujour for their advices in wheat culture and the management of the experimental site. We thank Tariq Shah for his contribution to the bibliographical search. We also thank Graines de Noé, especially Hélène Montaz, for providing the seeds of ancient varieties, as well as RAGT Semences, Secobra, Saaten Union and Limagrain for providing the modern varieties.

## AUTHOR CONTRIBUTIONS

MB conceived the research, MB wrote the paper, CD, EP and MB carried out the experiment, SJ dealt with sample preparation for sequencing and bioinformatics analysis, SJ did the statistical analyses and edited the figures and tables, DW, LC and TR did the mycorrhiza observations, SJ, TR, LC and DW edited and commented the paper.

