## Supporting Data for "Changes in wheat rhizosphere microbiota in response to chemical inputs, plant genotype and phenotypic plasticity" (from another version of this preprint: version 1, 2021-05-07)

-

[*]Corresponding author

**Running title**: Phenotypic plasticity of plant-microbiota interactions

**Figure S1:** Sequencing rarefaction curves of the raw samples. Curves are representing the microbial OTU detection rates as a function of recruited amplicon sequences, for the bacterial (A) and the fungal profiles (B), respectively. The dotted line indicates the rarefaction level that was chosen to randomly re-sample all profiles evenly to normalize the analysis. Curves marked in red are indicating samples that did not pass the quality control check and were excluded (See Table S1).

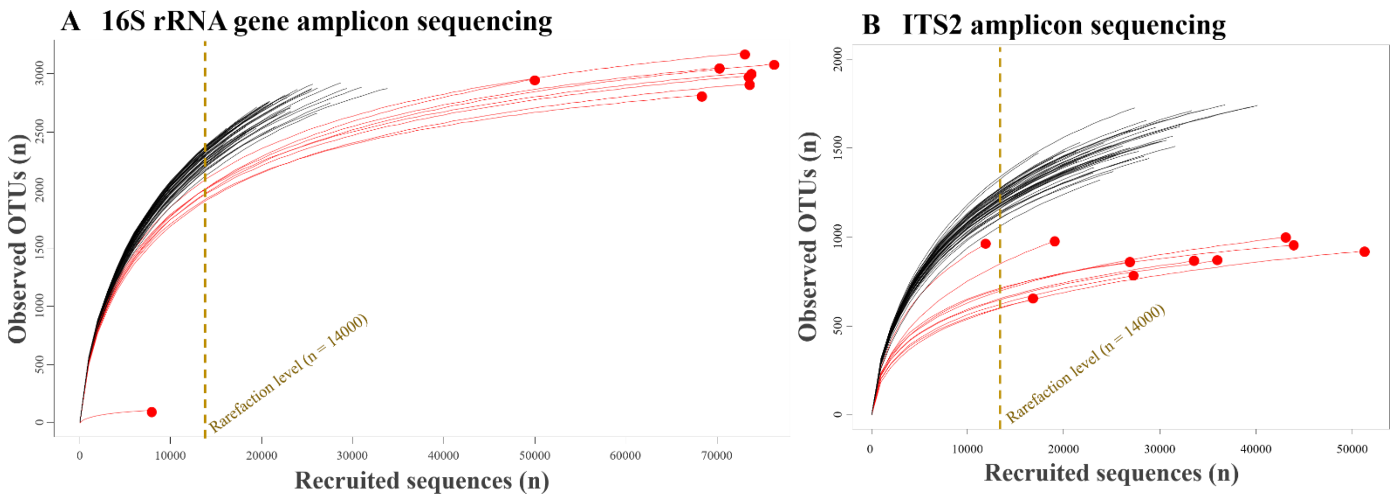

**Table S1:** Summary table of sequencing samples generated

| Amplicon_type | Sample | Variety | Inputs | Breed | Bloc | Sequences | Quality check |
|---|---|---|---|---|---|---|---|
| 16S rRNA gene | A_N_R1 | Alixan | w | modern | Bloc1 | 73639 | failed (column amplification issue) |
| 16S rRNA gene | A_N_R2 | Alixan | w | modern | Bloc2 | 16382 | passed |
| 16S rRNA gene | A_N_R3 | Alixan | w | modern | Bloc3 | 16131 | passed |
| 16S rRNA gene | A_SN_R1 | Alixan | w/o | modern | Bloc1 | 73092 | failed (column amplification issue) |
| 16S rRNA gene | A_SN_R2 | Alixan | w/o | modern | Bloc2 | 33815 | passed |
| 16S rRNA gene | A_SN_R3 | Alixan | w/o | modern | Bloc3 | 22629 | passed |
| 16S rRNA gene | AL_N_R1 | Alauda | w | ancient | Bloc1 | 7344 | failed (not enough sequences and outlier profil) |
| 16S rRNA gene | AL_N_R2 | Alauda | w | ancient | Bloc2 | 15748 | passed |
| 16S rRNA gene | AL_N_R3 | Alauda | w | ancient | Bloc3 | 17572 | passed |
| 16S rRNA gene | AL_SN_R1 | Alauda | w/o | ancient | Bloc1 | 23091 | passed |
| 16S rRNA gene | AL_SN_R2 | Alauda | w/o | ancient | Bloc2 | 29298 | passed |
| 16S rRNA gene | AL_SN_R3 | Alauda | w/o | ancient | Bloc3 | 15233 | passed |
| 16S rRNA gene | AR_N_R1 | Automne_Rouge | w | ancient | Bloc1 | 28594 | passed |
| 16S rRNA gene | AR_N_R2 | Automne_Rouge | w | ancient | Bloc2 | 14415 | passed |
| 16S rRNA gene | AR_N_R3 | Automne_Rouge | w | ancient | Bloc3 | 21209 | passed |
| 16S rRNA gene | AR_SN_R1 | Automne_Rouge | w/o | ancient | Bloc1 | 25452 | passed |
| 16S rRNA gene | AR_SN_R2 | Automne_Rouge | w/o | ancient | Bloc2 | 15723 | passed |
| 16S rRNA gene | AR_SN_R3 | Automne_Rouge | w/o | ancient | Bloc3 | 27576 | passed |
| 16S rRNA gene | BM_N_R1 | Barbu_Maconnais | w | ancient | Bloc1 | 18674 | passed |
| 16S rRNA gene | BM_N_R2 | Barbu_Maconnais | w | ancient | Bloc2 | 20874 | passed |
| 16S rRNA gene | BM_N_R3 | Barbu_Maconnais | w | ancient | Bloc3 | 21719 | passed |
| 16S rRNA gene | BM_SN_R1 | Barbu_Maconnais | w/o | ancient | Bloc1 | 18220 | passed |
| 16S rRNA gene | BM_SN_R2 | Barbu_Maconnais | w/o | ancient | Bloc2 | 23186 | passed |
| 16S rRNA gene | BM_SN_R3 | Barbu_Maconnais | w/o | ancient | Bloc3 | 23926 | passed |
| 16S rRNA gene | BS_N_R1 | Ble_Saone | w | ancient | Bloc1 | 30975 | passed |
| 16S rRNA gene | BS_N_R2 | Ble_Saone | w | ancient | Bloc2 | 17557 | passed |
| 16S rRNA gene | BS_N_R3 | Ble_Saone | w | ancient | Bloc3 | 19392 | passed |
| 16S rRNA gene | BS_SN_R1 | Ble_Saone | w/o | ancient | Bloc1 | 20842 | passed |
| 16S rRNA gene | BS_SN_R2 | Ble_Saone | w/o | ancient | Bloc2 | 18861 | passed |
| 16S rRNA gene | BS_SN_R3 | Ble_Saone | w/o | ancient | Bloc3 | 22992 | passed |
| 16S rRNA gene | D_N_R1 | Descartes | w | modern | Bloc1 | 25593 | passed |
| 16S rRNA gene | D_N_R2 | Descartes | w | modern | Bloc2 | 18705 | passed |
| 16S rRNA gene | D_N_R3 | Descartes | w | modern | Bloc3 | 19172 | passed |
| 16S rRNA gene | D_SN_R1 | Descartes | w/o | modern | Bloc1 | 19508 | passed |
| 16S rRNA gene | D_SN_R2 | Descartes | w/o | modern | Bloc2 | 16713 | passed |
| 16S rRNA gene | D_SN_R3 | Descartes | w/o | modern | Bloc3 | 18214 | passed |
| 16S rRNA gene | EN_N_R1 | Engrain | w | ancient | Bloc1 | 70433 | failed (column amplification issue) |
| 16S rRNA gene | EN_N_R2 | Engrain | w | ancient | Bloc2 | 21450 | passed |
| 16S rRNA gene | EN_N_R3 | Engrain | w | ancient | Bloc3 | 26074 | passed |
| 16S rRNA gene | EN_SN_R1 | Engrain | w/o | ancient | Bloc1 | 68322 | failed (column amplification issue) |
| 16S rRNA gene | EN_SN_R2 | Engrain | w/o | ancient | Bloc2 | 19747 | passed |
| 16S rRNA gene | EN_SN_R3 | Engrain | w/o | ancient | Bloc3 | 22500 | passed |
| 16S rRNA gene | R_N_R1 | Rubisco | w | modern | Bloc1 | 26116 | passed |
| 16S rRNA gene | R_N_R2 | Rubisco | w | modern | Bloc2 | 22675 | passed |
| 16S rRNA gene | R_N_R3 | Rubisco | w | modern | Bloc3 | 25487 | passed |
| 16S rRNA gene | R_SN_R1 | Rubisco | w/o | modern | Bloc1 | 50006 | failed (column amplification issue) |
| 16S rRNA gene | R_SN_R2 | Rubisco | w/o | modern | Bloc2 | 18990 | passed |
| 16S rRNA gene | R_SN_R3 | Rubisco | w/o | modern | Bloc3 | 23221 | passed |
| 16S rRNA gene | S_N_R1 | Sherlock | w | modern | Bloc1 | 18433 | passed |
| 16S rRNA gene | S_N_R2 | Sherlock | w | modern | Bloc2 | 18202 | passed |
| 16S rRNA gene | S_N_R3 | Sherlock | w | modern | Bloc3 | 19472 | passed |
| 16S rRNA gene | S_SN_R1 | Sherlock | w | modern | Bloc1 | 73883 | failed (column amplification issue) |
| 16S rRNA gene | S_SN_R2 | Sherlock | w/o | modern | Bloc2 | 22776 | passed |
| 16S rRNA gene | S_SN_R3 | Sherlock | w/o | modern | Bloc3 | 20734 | passed |
| 16S rRNA gene | T_N_R1 | Tulip | w | modern | Bloc1 | 73634 | failed (column amplification issue) |
| 16S rRNA gene | T_N_R2 | Tulip | w | modern | Bloc2 | 19489 | passed |
| 16S rRNA gene | T_N_R3 | Tulip | w | modern | Bloc3 | 16671 | passed |
| 16S rRNA gene | T_SN_R1 | Tulip | w/o | modern | Bloc1 | 82542 | failed (column amplification issue) |
| 16S rRNA gene | T_SN_R2 | Tulip | w/o | modern | Bloc2 | 20571 | passed |
| 16S rRNA gene | T_SN_R3 | Tulip | w/o | modern | Bloc3 | 16115 | passed |
| ITS2 fragment | A_N_R1 | Alixan | w | modern | Bloc1 | 27133 | failed (column amplification issue) |
| ITS2 fragment | A_N_R2 | Alixan | w | modern | Bloc2 | 21274 | passed |
| ITS2 fragment | A_N_R3 | Alixan | w | modern | Bloc3 | 21458 | passed |
| ITS2 fragment | A_SN_R1 | Alixan | w/o | modern | Bloc1 | 43298 | failed (column amplification issue) |

| | | | | | | | |
|---|---|---|---|---|---|---|---|
| ITS2 fragment | A_SN_R2 | Alixan | w/o | modern | Bloc2 | 34673 | passed |
| ITS2 fragment | A_SN_R3 | Alixan | w/o | modern | Bloc3 | 26886 | passed |
| ITS2 fragment | AL_N_R1 | Alauda | w | ancient | Bloc2 | 11942 | failed (not enough sequences and outlier profil) |
| ITS2 fragment | AL_N_R2 | Alauda | w | ancient | Bloc2 | 20081 | passed |
| ITS2 fragment | AL_N_R3 | Alauda | w | ancient | Bloc3 | 23782 | passed |
| ITS2 fragment | AL_SN_R1 | Alauda | w/o | ancient | Bloc1 | 25915 | passed |
| ITS2 fragment | AL_SN_R2 | Alauda | w/o | ancient | Bloc2 | 31331 | passed |
| ITS2 fragment | AL_SN_R3 | Alauda | w/o | ancient | Bloc3 | 17310 | passed |
| ITS2 fragment | AR_N_R1 | Automne_Rouge | w | ancient | Bloc1 | 28591 | passed |
| ITS2 fragment | AR_N_R2 | Automne_Rouge | w | ancient | Bloc2 | 19967 | passed |
| ITS2 fragment | AR_N_R3 | Automne_Rouge | w | ancient | Bloc3 | 29867 | passed |
| ITS2 fragment | AR_SN_R1 | Automne_Rouge | w/o | ancient | Bloc1 | 25452 | passed |
| ITS2 fragment | AR_SN_R2 | Automne_Rouge | w/o | ancient | Bloc1 | 17325 | passed |
| ITS2 fragment | AR_SN_R3 | Automne_Rouge | w/o | ancient | Bloc2 | 22695 | passed |
| ITS2 fragment | BM_N_R1 | Barbu_Maconnais | w | ancient | Bloc3 | 21280 | passed |
| ITS2 fragment | BM_N_R2 | Barbu_Maconnais | w | ancient | Bloc1 | 23114 | passed |
| ITS2 fragment | BM_N_R3 | Barbu_Maconnais | w | ancient | Bloc2 | 23057 | passed |
| ITS2 fragment | BM_SN_R1 | Barbu_Maconnais | w/o | ancient | Bloc3 | 22954 | passed |
| ITS2 fragment | BM_SN_R2 | Barbu_Maconnais | w/o | ancient | Bloc1 | 26947 | passed |
| ITS2 fragment | BM_SN_R3 | Barbu_Maconnais | w/o | ancient | Bloc2 | 33312 | passed |
| ITS2 fragment | BS_N_R1 | Ble_Saone | w | ancient | Bloc3 | 25791 | passed |
| ITS2 fragment | BS_N_R2 | Ble_Saone | w | ancient | Bloc1 | 25141 | passed |
| ITS2 fragment | BS_N_R3 | Ble_Saone | w | ancient | Bloc2 | 19233 | failed (not enough sequences and outlier profil) |
| ITS2 fragment | BS_SN_R1 | Ble_Saone | w/o | ancient | Bloc1 | 14658 | passed |
| ITS2 fragment | BS_SN_R2 | Ble_Saone | w/o | ancient | Bloc3 | 32019 | passed |
| ITS2 fragment | BS_SN_R3 | Ble_Saone | w/o | ancient | Bloc1 | 36741 | passed |
| ITS2 fragment | D_N_R1 | Descartes | w | modern | Bloc2 | 26156 | passed |
| ITS2 fragment | D_N_R2 | Descartes | w | modern | Bloc3 | 19217 | passed |
| ITS2 fragment | D_N_R3 | Descartes | w | modern | Bloc1 | 21503 | passed |
| ITS2 fragment | D_SN_R1 | Descartes | w/o | modern | Bloc2 | 19211 | passed |
| ITS2 fragment | D_SN_R2 | Descartes | w/o | modern | Bloc3 | 28312 | passed |
| ITS2 fragment | D_SN_R3 | Descartes | w/o | modern | Bloc1 | 24980 | passed |
| ITS2 fragment | EN_N_R1 | Engrain | w | ancient | Bloc1 | 36024 | failed (column amplification issue) |
| ITS2 fragment | EN_N_R2 | Engrain | w | ancient | Bloc2 | 30149 | passed |
| ITS2 fragment | EN_N_R3 | Engrain | w | ancient | Bloc3 | 25457 | passed |
| ITS2 fragment | EN_SN_R1 | Engrain | w/o | ancient | Bloc1 | 27296 | failed (column amplification issue) |
| ITS2 fragment | EN_SN_R2 | Engrain | w/o | ancient | Bloc2 | 29586 | passed |
| ITS2 fragment | EN_SN_R3 | Engrain | w/o | ancient | Bloc3 | 40208 | passed |
| ITS2 fragment | R_N_R1 | Rubisco | w | modern | Bloc2 | 14441 | passed |
| ITS2 fragment | R_N_R2 | Rubisco | w | modern | Bloc3 | 23207 | passed |
| ITS2 fragment | R_N_R3 | Rubisco | w | modern | Bloc1 | 27386 | passed |
| ITS2 fragment | R_SN_R1 | Rubisco | w/o | modern | Bloc1 | 43986 | failed (column amplification issue) |
| ITS2 fragment | R_SN_R2 | Rubisco | w/o | modern | Bloc2 | 27797 | passed |
| ITS2 fragment | R_SN_R3 | Rubisco | w/o | modern | Bloc3 | 21413 | passed |
| ITS2 fragment | S_N_R1 | Sherlock | w | modern | Bloc1 | 21736 | passed |
| ITS2 fragment | S_N_R2 | Sherlock | w | modern | Bloc2 | 26770 | passed |
| ITS2 fragment | S_N_R3 | Sherlock | w | modern | Bloc3 | 25136 | passed |
| ITS2 fragment | S_SN_R1 | Sherlock | w/o | modern | Bloc1 | 51516 | failed (column amplification issue) |
| ITS2 fragment | S_SN_R2 | Sherlock | w/o | modern | Bloc2 | 39700 | passed |
| ITS2 fragment | S_SN_R3 | Sherlock | w/o | modern | Bloc3 | 28833 | passed |
| ITS2 fragment | T_N_R1 | Tulip | w | modern | Bloc1 | 16999 | failed (column amplification issue) |
| ITS2 fragment | T_N_R2 | Tulip | w | modern | Bloc2 | 31623 | passed |
| ITS2 fragment | T_N_R3 | Tulip | w | modern | Bloc3 | 24181 | passed |
| ITS2 fragment | T_SN_R1 | Tulip | w/o | modern | Bloc1 | 33749 | failed (column amplification issue) |
| ITS2 fragment | T_SN_R2 | Tulip | w/o | modern | Bloc2 | 30717 | passed |
| ITS2 fragment | T_SN_R3 | Tulip | w/o | modern | Bloc3 | 18837 | passed |

**Table S2:** ANOVA models of the normally distributed bacterial alpha-diversity indices. Df = degree of freedom, SoS = Sum of Squares, F = Fischer index, Var. SoS = percentage of variance based on the sum of squares, p = p-value, Signif = significance, ˙, $p < 0.10$; *, $p < 0.05$; **, $p < 0.01$; ***, $p < 0.001$.

| Index | Factors | Df | SoS | F | Var. SoS (%) | *p* | Signif. |
|---|---|---|---|---|---|---|---|
| Richness | inputs | 1 | 149 | 0.06 | 0.06 | 0.808 | |
| | breeding type | 1 | 36673 | 14.81 | 15.57 | 6.0E-05 | *** |
| | block | 2 | 212 | 0.04 | 0.09 | 0.958 | |
| | inputs:breeding type | 1 | 18230 | 7.36 | 7.74 | 0.011 | * |
| | inputs:breeding type:variety | 16 | 108531 | 2.7 | 46.07 | 0.009 | ** |
| | Residuals | 29 | 71803 | | 30.48 | | |
| Chao1 | inputs | 1 | 23478 | 1.69 | 2.08 | 0.204 | |
| | breeding type | 1 | 77711 | 5.59 | 6.89 | 0.025 | * |
| | block | 2 | 12360 | 0.44 | 1.10 | 0.646 | |
| | inputs:breeding type | 1 | 47461 | 3.41 | 4.21 | 0.075 | . |
| | inputs:breeding type:variety | 16 | 562809 | 2.53 | 49.93 | 0.015 | * |
| | residuals | 29 | 403443 | | 35.79 | | |
| ACE | inputs | 1 | 2799 | 0.21 | 0.27 | 0.653 | |
| | breeding type | 1 | 60056 | 4.44 | 5.87 | 0.044 | * |
| | block | 2 | 11838 | 0.44 | 1.16 | 0.650 | |
| | inputs:breeding type | 1 | 69730 | 5.15 | 6.81 | 0.031 | * |
| | inputs:breeding type:variety | 16 | 486652 | 2.25 | 47.54 | 0.028 | * |
| | residuals | 29 | 392616 | | 38.35 | | |

**Table S3:** ANOVA models of the normally distributed fungal alpha-diversity indices. Df = degree of freedom, SoS = Sum of Squares, F = Fischer index, Var. SoS = percentage of variance based on the sum of squares, p = p-value, Signif = significance, ˙, p < 0.10; *, p < 0.05; **, p < 0.01; ***, p < 0.001.

| Index | Factors | Df | SoS | F | Var. SoS (%) | *p* | Signif. |
|---|---|---|---|---|---|---|---|
| Richness | inputs | 1 | 4287 | 1.00 | 2.61 | 0.325 | |
| | breeding type | 1 | 2 | 1.0E-4 | 1.0E-4 | 0.982 | |
| | block | 2 | 7905 | 0.92 | 4.81 | 0.409 | |
| | inputs:breeding type | 1 | 12167 | 2.85 | 7.40 | 0.103 | |
| | inputs:breeding type:variety | 16 | 20404 | 0.30 | 12.40 | 0.993 | |
| | residuals | 28 | 119756 | | 72.80 | | |
| Chao1 | inputs | 1 | 59 | 0.01 | 0.01 | 0.947 | |
| | breeding type | 1 | 154 | 0.01 | 0.03 | 0.914 | |
| | block | 2 | 30046 | 1.15 | 5.7 | 0.332 | |
| | inputs:breeding type | 1 | 73602 | 5.62 | 13.9 | 0.025 | * |
| | inputs:breeding type:variety | 16 | 58994 | 0.28 | 11.1 | 0.995 | |
| | Residuals | 28 | 366469 | | 69.2 | | |
| ACE | inputs | 1 | 809 | 0.05 | 0.1 | 0.829 | |
| | breeding type | 1 | 69 | 0.00 | 0.0 | 0.950 | |
| | block | 2 | 23011 | 0.68 | 3.7 | 0.516 | |
| | inputs:breeding type | 1 | 52634 | 3.10 | 8.4 | 0.089 | . |
| | inputs:breeding type:variety | 16 | 76652 | 0.28 | 12.2 | 0.995 | |
| | residuals | 28 | 475909 | | 75.7 | | |
| Shannon index | inputs | 1 | 0.038 | 4.46 | 6.7 | 0.044 | * |
| | breeding type | 1 | 0.008 | 0.90 | 1.4 | 0.351 | |
| | block | 2 | 0.111 | 6.57 | 19.8 | 0.005 | ** |
| | inputs:breeding type | 1 | 0.026 | 3.04 | 4.6 | 0.093 | . |
| | inputs:breeding type:variety | 16 | 0.142 | 1.05 | 25.3 | 0.441 | |
| | residuals | 28 | 0.237 | | 42.2 | | |
| Equitability | inputs | 1 | 5.0E-04 | 3.60 | 5.4 | 0.068 | . |
| | breeding type | 1 | 2.0E-04 | 1.12 | 1.7 | 0.298 | |
| | block | 2 | 0.002 | 5.99 | 18.0 | 0.007 | ** |
| | inputs:breeding type | 1 | 2.0E-04 | 1.41 | 2.1 | 0.245 | |
| | inputs:breeding type:variety | 16 | 0.003 | 1.28 | 30.7 | 0.278 | |
| | residuals | 28 | 0.004 | | 42.1 | | |

**Table S4:** ANOVA models of the normally distributed mycorrhiza indices. Df = degree of freedom, SoS = Sum of Squares, F = Fischer index, Var. SoS = percentage of variance based on the sum of squares, p = p-value, Signif = significance, ˙, *p* < 0.10; *, *p* < 0.05; **, *p* < 0.01; ***, *p* < 0.001.

| Index | Factors | Df | SoS | F | Var. SoS (%) | *p* | Signif. |
|---|---|---|---|---|---|---|---|
| | inputs | 1 | 193 | 5.03 | 2.65 | 0.027 | * |
| | breeding | 1 | 311 | 8.08 | 4.26 | 0.005 | ** |
| | serie | 1 | 997 | 25.93 | 13.67 | 1.7E-7 | *** |
| M | bloc | 2 | 301 | 3.92 | 4.13 | 0.023 | * |
| | inputs:breeding | 1 | 38 | 0.99 | 0.52 | 0.321 | |
| | inputs:breeding:variety | 16 | 1724 | 2.80 | 23.63 | 9.6E-5 | *** |
| | residuals | 97 | 3731 | | 51.14 | | |
| | inputs | 1 | 199 | 4.06 | 2.59 | 0.047 | * |
| | breeding | 1 | 349 | 7.12 | 4.53 | 0.009 | ** |
| | serie | 1 | 341 | 6.97 | 4.43 | 0.010 | ** |
| m | bloc | 2 | 297 | 3.03 | 3.86 | 0.053 | . |
| | inputs:breeding | 1 | 30 | 0.62 | 0.39 | 0.433 | |
| | inputs:breeding:variety | 16 | 1729 | 2.21 | 22.46 | 0.009 | ** |
| | residuals | 97 | 4753 | | 61.7 | | |
| | inputs | 1 | 45 | 0.18 | 0.12 | 0.673 | |
| | breeding | 1 | 1434 | 5.75 | 3.82 | 0.018 | * |
| | serie | 1 | 3399 | 13.62 | 9.06 | 3.7E-5 | *** |
| a | bloc | 2 | 486 | 0.97 | 1.29 | 0.381 | |
| | inputs:breeding | 1 | 545 | 2.19 | 1.45 | 0.143 | |
| | inputs:breeding:variety | 16 | 7421 | 1.86 | 19.77 | 0.034 | * |
| | residuals | 97 | 24200 | | 64.48 | | |
